# A reassessment of DNA immunoprecipitation-based genomic profiling

**DOI:** 10.1101/224279

**Authors:** Antonio Lentini, Cathrine Lagerwall, Svante Vikingsson, Heidi K. Mjoseng, Karolos Douvlataniotis, Hartmut Vogt, Henrik Green, Richard R. Meehan, Mikael Benson, Colm E. Nestor

## Abstract

DNA immunoprecipitation sequencing (DIP-seq) is a common enrichment method for profiling DNA modifications in mammalian genomes. However, DIP-seq profiles often exhibit significant variation between independent studies of the same genome and from profiles obtained by alternative methods. Here we show that these differences are primarily due to intrinsic affinity of IgG for short unmodified DNA repeats. This pervasive experimental error accounts for 50 – 99% of regions identified as ‘enriched’ for DNA modifications in DIP-seq data. Correction of this error profoundly alters DNA modification profiles for numerous cell types, including mouse embryonic stem cells, and subsequently reveals novel associations between DNA modifications, chromatin modifications and biological processes. We propose new methodological guidelines that minimize the impact of these errors on future DIP-seq experiments and allow new insights to be made from the wealth of existing DIP-seq data.

---

Canonical DNA methylation in mammals involves the covalent attachment of a methyl group to cytosine to form 5-methylcytosine (5mC). The ability to establish and maintain DNA methylation patterns is essential for normal development in mammals, and aberrant DNA methylation is observed in numerous diseases, including all forms of cancer^1^. Comprehensive mapping of DNA methylation (5-methylcytosine, 5mC) in multiple species has been critical to establishing the relevance of methylation dynamics in gene regulation and chromatin organization^2–4^. An effective method of generating genome-wide 5mC profiles couples antibody-based enrichment of methylated DNA fragments (MeDIP) with hybridization to DNA micro-arrays (MeDIP-chip) or high-throughput sequencing (MeDIP-seq)^5, 6^. Subsequent comparisons with nucleotide resolution bisulphite sequencing (BS) techniques produced broadly correlative DNA methylation data^6, 7^. Unlike BS, the MeDIP-seq information is not contained in the read sequence itself, but in the enrichment or depletion of sequencing reads that map to specific regions of the genome^7, 8^. Consequently, appropriate control samples are required, which typically correspond to the input genomic DNA before enrichment. The low cost of DIP-seq initially made it the method of choice in studies involving large numbers of samples. Subsequently the application of the DIP-seq technique has been extended to chart the genomic location of additional DNA modifications (5-hydroxymethylcytosine, 5hmC; 5-Lentini *et al.*formylcytosine, 5fC; 5-carboxycytosine, 5caC; and 6-methyladenosine, 6mA) as their corresponding antibodies became available, elucidating their roles in the process of DNA methylation remodeling and gene regulation^9-16^. Interestingly, verification of DIP profiles by independent methods revealed several problems with the DIP-seq approach, including preferential enrichment of low CG content regions by the 5mC antibody^17^ and enrichment of highly modified regions by the 5hmC antibody^18^. In addition, we and others recently reported an enrichment of short tandem repeat (STR) sequences in hMeDIP assays^19, 20^. However, the origin of STR enrichment and the scale of its impact on DIP-seq data remained unknown.

Here we performed a systematic analysis of DIP-seq profiles generated with antibodies against multiple DNA modifications.We demonstrate that highly specific off-target binding to unmodified repetitive sequences is not limited to 5hmC antibodies but is an inherent technical error observed in all DIP-seq studies, irrespective of the target DNA modification, cell-type or organism. We reveal that between 50% – 99% of enriched regions in DIP-Seq data are false positives, the removal of which markedly affects our perception of methylation dynamics in mammals; altering the associations between DNA methylation and other genomic and epigenomic features. In addition to inherent errors in DIP-seq, we also observed that contamination of mammalian samples with 6mA containing bacterial DNA may account for the conflicting findings relating to the location and abundance of 6mA in mammalian genomes. Finally, we detail adjustments to existing DNA immunoprecipitation protocols and suggest novel computational approaches that will minimize the impact of these errors on future DIP-seq experiments and allow new insights to be gained from the wealth of existing DIP-seq data.

## RESULTS

### IgG has an intrinsic affinity for short tandem repeats in mammalian DNA

To simplify comparison of DIP-seq results from separate studies we used a uniform computational pipeline (see **online methods**) to analyze published DIP-seq profiles of 5mC, 5hmC, 5fC and 5caC (hereby referred to as ‘5modC’) in mouse embryonic stem cells (mESCs). The sensitivity and specificity of 5modC antibodies used in DIP-Seq is well established,with limited to undetectable cross-reactivity observed in dot-blot and ELISA assays^15, 19, 20^ (**Supplementary Fig.1a, b**). All analyzed datasets and their relationship to figures is outlined in **Supplementary Table 1**. This approach revealed a striking enrichment at short tandem repeats (STRs) in all 5modC DIP-seq datasets (**Fig. 1a**). Surprisingly, near identical enrichment patterns at STRs were observed in mESC DIP-seq generated with a non-specific mouse IgG antibody (**Fig. 1a**). The intersection of regions enriched for all 5modC showed a 5.8 fold higher enrichment for IgG compared to non-enriched DNA (Input; *P*=5.03x10^-5^,T-test) whereas nonintersecting regions showed no difference (**Supplementary Fig. 1c**), suggesting that a proportion of the 5modC signal may be due to off-target binding of the antibodies. Indeed, genome-wide IgG enrichment could explain up to 55% of all 5modC DIP-seq enriched loci in mESCs whereas Input explained a maximum of 3% of enriched regions (**Supplementary Fig. 1d**). Significantly, overlapping 5mC, 5hmC and IgG regions were depleted of CpG dinucleotides compared to regions not overlapping IgG (**Supplementary Fig. 1e**). Although non-CpG methylation is known to occur in mESCs ^21, 22^, analysis of whole-genome bisulfite sequencing data_21_ confirmed that CpHs in these regions were primarily unmethylated (median Lentini methylated C_p_H_s_ = 0 and 3 for IgG and 5mC regions, respectively) (**Supplementary Fig. 1f**) suggesting that all antibodies were binding highly specific regions of unmodified DNA during DIP experiments. We verified this by analyzing published DIP-seq data from Dnmt triple knockout (TKO) mESCs^23^ that lack DNA methyltransferase activity and revealed that the 5hmC antibody enriched similar regions to that of the IgG control (**Fig. 1b** and **Supplementary Fig. 1g**). We confirmed depletion of both 5mC and 5hmC in TKO compared to wild-type (WT)mESC DNA using mass spectrometry (**Fig. 1c**), verifying that the DIP-seq signals observed in TKO cells were independent of 5modC status. 5hmC–DIP followed by qPCR confirmed the enrichment of STRs in TKO mESCs lacking 5hmC (**Fig. 1d,e**). Significantly, 5hmC profiles generated from an independent, non-antibody based 5hmC enrichment technique^24^ (5hmC-Seal) showed no enrichment over IgG regions (**Fig. 1f** further implicating off-target binding of STRs by antibodies during DIP-seq

**Figure 1.**
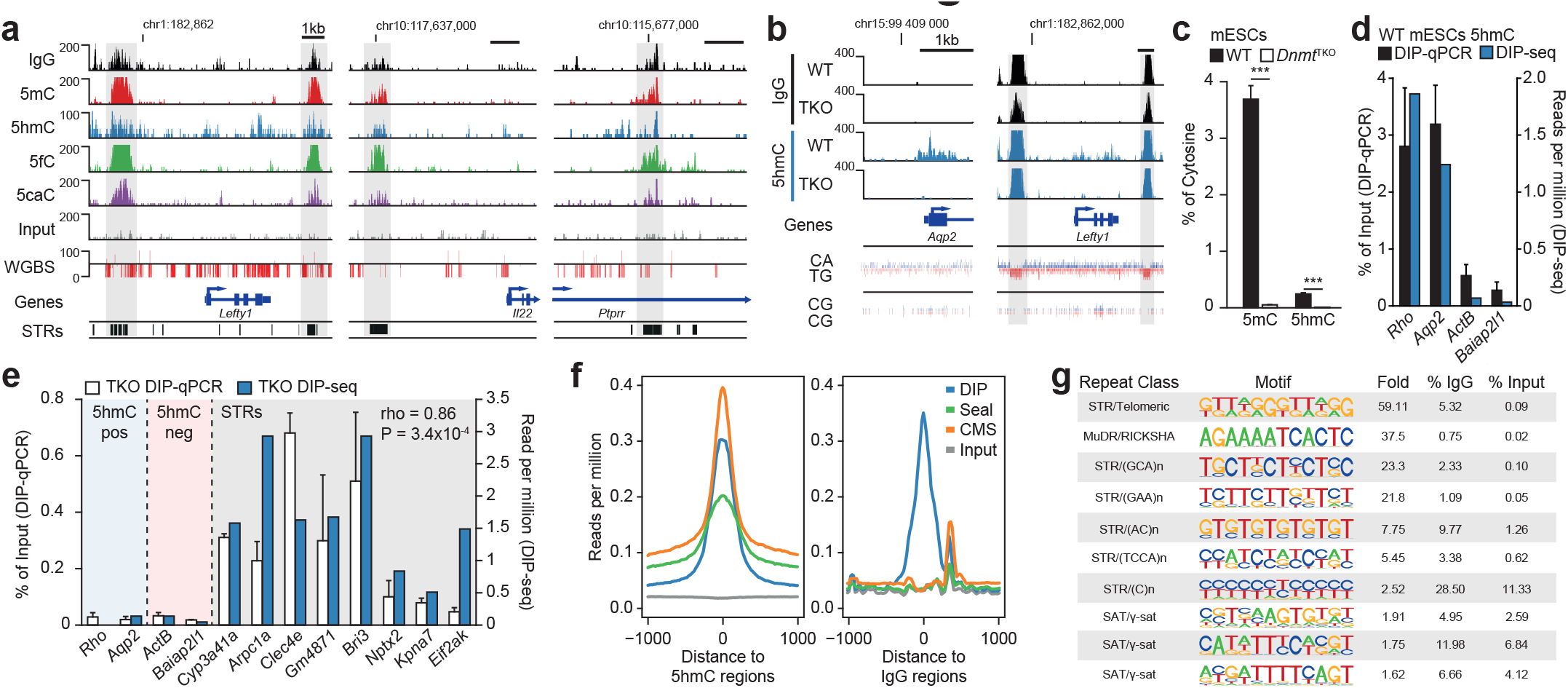
Characterization of off-target antibody binding in DIP-seq.(**a**) Signal track in mESCsshowing similar enrichment between 5modC and IgG DIP-seq samples over repetitive regions.WGBS, whole-genome bisulfite sequencing; STRs, short tandem repeats. (**b**) Signal track of5hmC and IgG DIP-seq in hypomethylated *Dnmt* triple knockout (TKO) or methylated wild—type (WT) mESCs over 5hmC-(left) or IgG enriched regions (right). (**c**) MassspectrometryLentini *et al.*quantification of 5mC and 5hmC in TKO and WT mECSs showing their depletion in TKO. Bars represent mean ± s.d of 3 biological replicates. ***P-value < 0.001, T-test. (**d**) DIP-qPCR measurement of 5hmC levels in *Dnmt* wild-type (WT) mESCs over positive and negativecontrol regions. Bars represent mean ± s.d of 3 biological replicates. (**e**) DIP-qPCR measurement using a 5hmCantibody in *Dnmt* triple knockout (TKO) mESCs showing off-target enrichment over repetitive regions. Bars represent mean ± s.d of 3 biological replicates, correlation calculated using Spearman correlation. STRs, short tandem repeats. (**f**) 5hmC enrichment in mESCs with different profiling techniques over 5hmC-(left) orIgG enriched regions (right) showing specific DIP-seq enrichment over IgG regions. (**g**) Motif enrichment for raw IgG reads compared toInput showing enrichment of repetitive sequences of short tandem repeat (STR) and satellite (SAT) classes of repeats.

To identify specific IgG-bound sequences, we screened the raw sequencing reads from three IgG DIP-seq samples in mESCs for overrepresented sequences, which revealed that between 30 and 60% of all reads were significantly enriched for repetitive motifs compared to Input (**Fig. 1g** and **Supplementary Table 2**), including the previously reported CA-repeats^19^. This suggested that IgG antibodies may have an innate binding capacity for single stranded repetitive DNA sequences. Interestingly, one of the most enriched motifs in IgG sequences reads was the 6-base TTAGGG sequence, suggesting that IgG also binds telomeric DNA which is susceptible to the formation of non-duplex (four-stranded quadruplex) structures^25^. Not only were IgG-DIP enriched for repetitive motifs, but the enriched IgG motifs were highly similar between samples (average Pearson *r* = 0.72) indicating that IgG binding is specific and reproducible (**Supplementary Table 2**). We observed similar repeat motifs in 5modC DIP-seq data from mESCs as well as a recently published study in mouse embryonic fibroblasts (MEFs)^26^ (r mESC = 0.75, r MEF = 0.68, **Supplementary Table 2**), showing that off-target binding of STRs in DIP-seq is not limited to mESCs and is highly sequence dependent. Indeed, the only antibody based profiling technique that did not show enrichment over IgG enriched regions was cytosine-5-methylenesulfonate (CMS)-seq^27^ (**Fig. 1f**), which involves bisulfite conversion of all unmodified cytosines to thymine before immunoprecipitation with the anti-CMS antibody. Consequently, all unmodified CA-repeats would be converted to TA-repeats.The lack of CA-repeat enrichment in CMS-Seq is thus strongly supportive of sequence-specific off-target binding of STRs by IgG antibodies. Taken together, our analyses indicates that native DNA immunoprecipitation libraries generated with multiple cytosine modification antibodies enriches for highly specific regions of unmodified repetitive DNA. This systematic error has resulted in extremely inaccurate attributions with respect to the genomic location of these modifications in mammals.

### IgG binding of DNA repeats explains the conflicting results of 6mA profiling in vertebrates

Next, we extended our analysis to a non-cytosine modification, 6-methyldeoxyadenosine (6mA), that is abundant in many bacteria and recently characterized in invertebrates^10, 28–30^. Its subsequent discovery in mammalian DNA has sparked an intense research effort to verify its location and characterize its function^13, 31^. To determine if 6mA DIP-seq studies have also been affected by off-target binding of STRs we compared profiles from primary mouse kidney cells^13^ to mESC IgG DIP-seq profiles. Again, 6mA profiles showed a clear enrichment at STRs and were highly similar to the mESC IgG profile (**Fig. 2a, b**). Analysis of additional public datasets revealed that 6mA DIP-seq data for *Danio rerio*^14^ and *Xenopus laevis^13^* also showed similar off-target enrichment for the same STR motifs observed in 5modC DIP-seq, whereas the 6mA rich genomes of *C. elegans^29^* and *E.coli^13^* showed no enrichment for these motifs (**Fig. 2c, d**). Inter-species differences in STR enrichment reflected the frequency of STRs in the genome of each species (**Fig. 2e**). Surprisingly, data from a recent study of 6mA-DIP in mESCs^31^ displayed different enrichment profiles from both mouse kidney 6mA- and mESC IgG-DIP-seq. Since the mESCs were cultured *in vitro* we wanted to exclude possible contamination of 6mA-rich bacteria such as *Mycoplasma*^32, 33^. To test this we mapped sequencing data to a combined genome index of *M.musculus, Mycosplasma sp.* and *E.coli* as a control (**see Online Methods**). This revealed that up to 15%of DIP-seq reads from the 6mA mESC study^31^ mapped to *Mycoplasma* whereas 24 samples from four other studies^13, 15, 23, 26^ mapped exclusively to *M.musculus* (**Fig. 2f**). Contamination of these samples may explain the earlier detection of 6mA in mammalian samples by mass spectrometry^31^ and the subsequent failure of more recent attempts to detect 6mA in mammalian DNA using ultrasensitive UHPLC-MS ^28, 34^.

**Figure 2.**
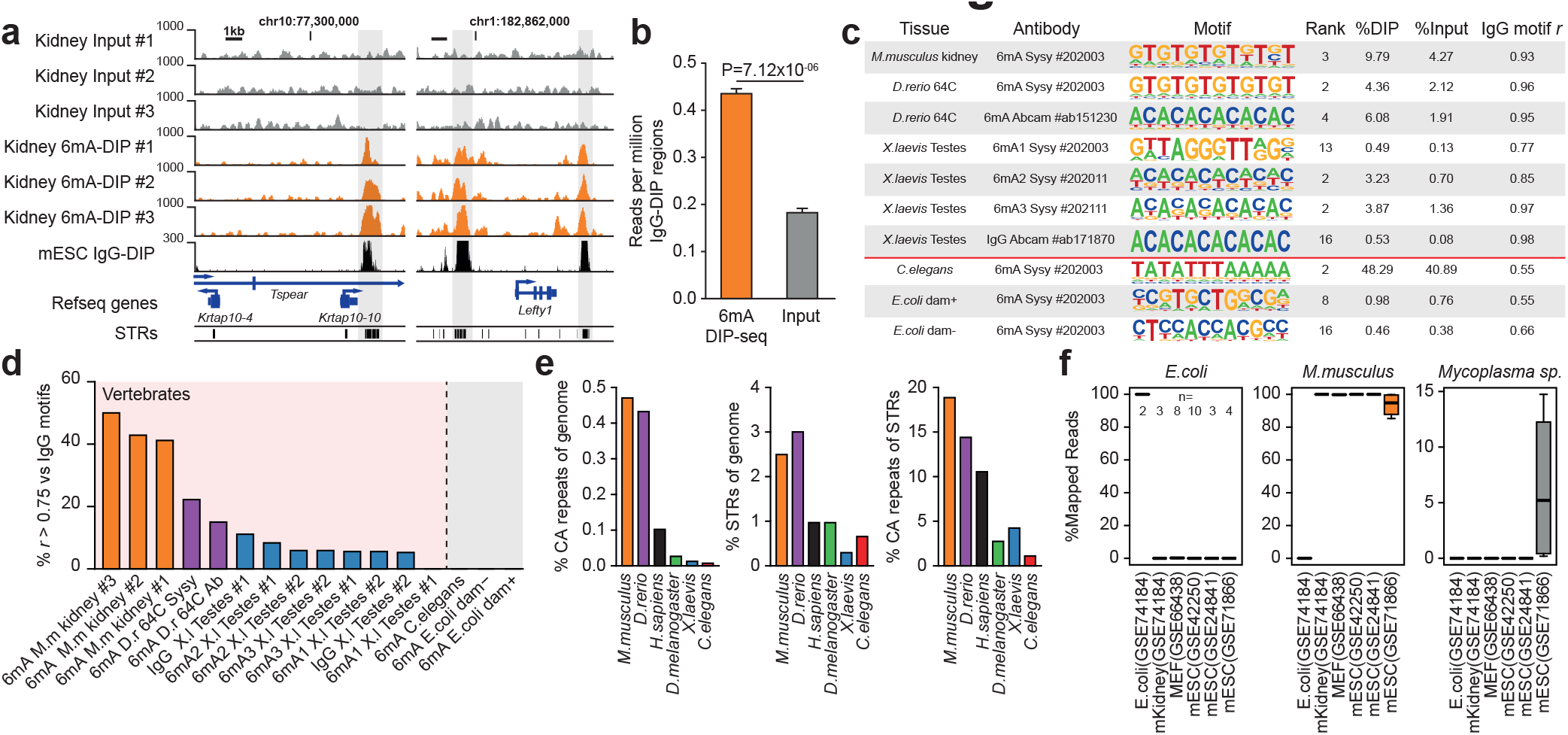
Characterization of similarities between 6mA and IgG DIP-seq in different species. (**a**) Signal track for Input and 6mA DIP-seq in mouse kidney cells and IgG DIP-seq in mESCs. (**b**) 6mA DIP-seq and Input enrichment over IgG enriched DIP-seq regions. Bars represent mean ± s.d of 3biological replicates. P—values calculated using T-test. STRs, short tandem repeats (**c**) Motif enrichment for raw 6mAor IgG DIP-seqreads compared to Input in multiple species. Motif with highest correlation to IgG motifs shown for each dataset. 6mA DIP-seq motifs in invertebrates show low correlation with IgG DIP-seq motifs whereas vertebrates show high correlation. (**d**) Barplot showing percent of motifs highly similar (*r* > 0.75) to IgG motifs for multiple species. (**e**) Proportion of short tandem repeats (STRs) in the genome of model organisms. STRs, short tandem repeats (**f**) Percentage reads mapping to mouse- or bacterial genomes. Boxplots represent median and top/bottom quartiles.

### Normalizing for off-target STR binding sharpens our view of epigenetic organization in mammals

Whereas the use of an appropriate IgG control sample would normalize for the effect of enrichment of unmodified STRs during DIP, the vast majority (> 96%) of published DIP-seq studies do not use an IgG control. To determine how STR binding in DIP-seq has affected our understanding of DNA methylation in mammals, we reanalyzed data from five independent studies of 5modC marks in mESCs^15, 19, 23, 35, 36^. First, we estimated the fraction of falsely enriched regions when using Input as a control, finding that up to 99% of enriched 5fC and 5caC, and approximately half of all 5hmC and 5mC regions could be considered false positives (**Fig. 3a**). In contrast, the mean percentage of falsely enriched regions was only 5.6% for all 5modC marks when using IgG as a control (**Fig. 3a**). These results suggested that the 5modC landscape in mammalian genomes has been greatly overestimated by DIP-seq (**Supplementary Fig. 2a, b**). Indeed, correcting for IgG not only reduced the number of enriched regions but also greatly increased the overlap with CMS and Seal profiling techniques (**Supplementary Fig. 2c**). Not surprisingly, the proportion of enriched repeat types was markedly altered when using Input or IgG controls in DIP-Seq, with STRs showing the greatest changes in enrichment (**Supplementary Fig.2d**). Interestingly,in addition to CA/GT repeats, AG/CT repeats were also highly enriched in Input controlled studies (**Supplementary Fig. 2e**).Off-target binding of AG rich motifs may explain the reported association of 6mA with specific AG-rich repeats (AGGG_N_)^13^ which was highly similar to those observed for all 5modC-DIP and IgG-DIP in mESCs (**Supplementary Table 2**).

**Figure 3.**
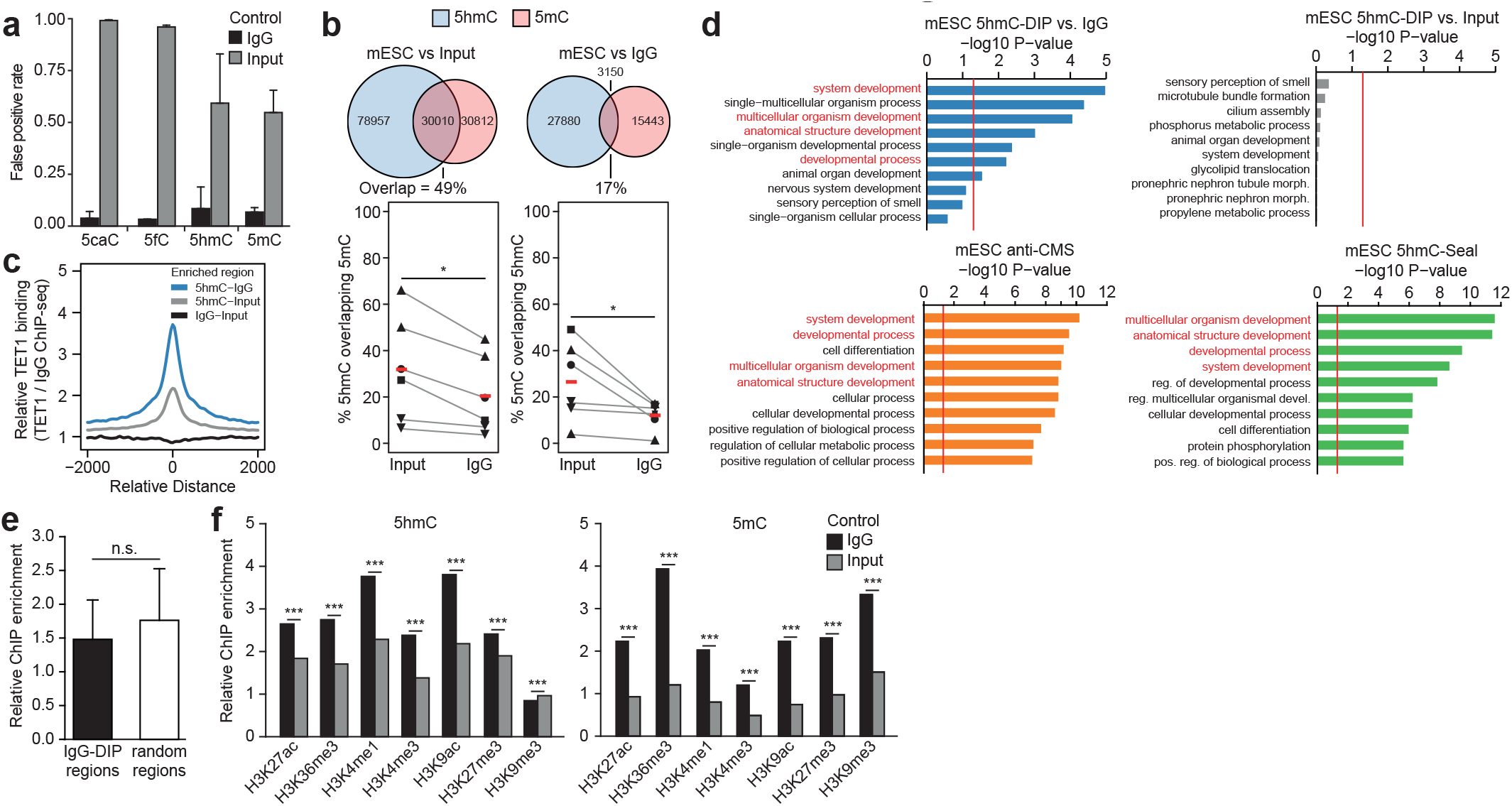
Biological impact of IgG correction. (**a**) Estimated false positive rate of enriched regions using IgG orInput ascontrol. Bars represent mean ± s.d of 2–7 biological replicates.(**b**) Overlap of 5hmC and 5mC regions using IgG or Input controls showing decreased overlapLentini *et al.*when using IgG controls. Venn diagram of 5mC and 5hmC overlap using IgG or Input controls (top). Below is a paired line plot of 5mC and 5hmC overlap using IgG or Input controls for multiple studies (indicated by symbols). Red markers indicate mean, *P<0.05, paired T-test. ▲ = ERP000570, ● = GSE31343, ◼ = GSE24841, ▼ = GSE42250. (**c**) TET1 binding over IgG- or 5hmC regions using IgG or Input controls. (**d**) GO term enrichment of biological processes for 5hmC enriched regions in mESCs using either IgG or Input as controls showing stronger enrichment for developmental terms when using IgG controls and highly similar results to GO terms for 5hmC-Seal and anti-CMS enriched regions. (**e**) Enrichment of ENCODE mESC histone ChIP-seq data relative to controls in regions enriched for IgG in DIP-seq or random regions of same size and chromosome shows no difference in ChIP enrichment. Bars represent mean ± s.d. of 26 biological replicates, P-values calculated using T-test. (**f**) Enrichment of ENCODE mESC histone ChIP-seq data for 5hmC- (left) or 5mC (right) enriched regions using IgG or Input as controls showing stronger association when using IgG controls. Data presented as mean (IgG) and bootstrap mean (Input) of 26 biological replicates, ***P<1e-5,bootstrap resampling.

Having shown that correction for STR-binding markedly altered the profiles of all 5modC in mESCs, we next investigated if these altered profiles impacted on their predicted roles in DNA de-methylation and gene regulation. Globally, 49%of 5mC-co-located with 5hmC enriched regions when using Input, whereas only 17% were coincident for both 5hmC and 5mC when using IgG (**Fig. 3b**). This suggested a more restricted role for 5hmC mediated DNA de-methylation in the reprogramming of the mESC epigenome, an assertion supported by the markedly improved association between 5hmC and TET protein occupancy in the mESC genome upon normalization to IgG (**Fig. 3c**). Significantly, removal of signals caused by STR-binding by normalization to IgG also altered the association of 5hmC with biological pathways from non-significant associations with unrelated processes including ‘cilia formation’, ‘smell perception’ and ‘phosphorus metabolism’ to highly significant associations with processes related to mammalian development and cell differentiation (**Fig. 3d, upper panels**). Significantly, the 5hmC-associated biological processes identified after correction for STR-binding were highly similar to those obtained with 5hmC-Seal and CMS-Seq, which do not enrich for unmodified repeats (**Fig. 3d, lower panels**). An improved association with developmental and differentiation related processes was also observed when the same correction was applied to mouse embryonic fibroblast cells **Supplementary Fig. 2f**).

Finally, histone ChIP-seq data in mESCs from ENCODE^37^ showed no difference in enrichment between IgG DIP-seq enriched regions and randomly sampled regions (**Fig. 3e**) suggesting that repeats found in intact chromatin structures are not bound by IgG, possibly due to their inability to form secondary structures. Again, using an IgG control significantly increased the association of 5hmC with permissive histone marks in mESCs^37^ whereas the association with heterochromatin (H3K9me3) decreased (**Fig. 3f**). For 5mC, the association with histone marks was also significantly increased, accentuating co-localization with H3K9me3 as well as H3K36me3 which together with 5mC is involved in mRNA splicing^38^ (**Fig. 3f**). Taken together, these findings further highlight the profound effect off-target STR binding has had on our understanding of the interrelationship between 5modC with mammalian chromatin signatures but also demonstrates the wealth of novel discoveries to be made by re-analysis of the vast body of published DIP-seq data.

## DISCUSSION

Our reanalysis of published DIP-seq data revealed that all commonly used DIP-seq antibodies bind unmodified short tandem repeat (STR) sequences. By analyzing DIP-seq data from mouse embryonic stem cells (mESCs) lacking both 5mC and 5hmC we confirmed that STR binding was modification independent;as non-specific IgG antibody generated profiles highly similar to that of 5mC and 5hmC antibodies. Consequently, only studies that have normalized DNA modification enrichment to an IgG control have corrected for off-target binding^15, 23^ (**Fig. 4**)^15, 23^. Unfortunately, greater than 96% of published DIP-seq studies do not include an IgG control. We showed that between 50 to 99% of enriched regions are due to off-target STR binding in 5modC DIP studies, with studies of low abundance modifications (i.e. 5caC & 5fC)having the highest false positive rates.Although our findings require the field to re-visit published DIP-seq data, they also provide an exciting opportunity to rapidly further our understanding of the dynamics of DNA methylation in mammalian biology by re-assessing the wealth of published DIP-seq data deposited in public databanks. Controlling for off-target IgG binding increased the signal to noise ratio in DIP-seq assays > 3-fold, allowing identification of more subtle alterations in modification levels. This also results in a significantly smaller and more distinct epigenomic landscape in mammalian cells, evidenced by a significantly reduced overlap between 5mC and 5hmC marked loci and a stronger association between 5modC and a variety of chromatin marks (**Fig. 3**).

**Figure 4.**
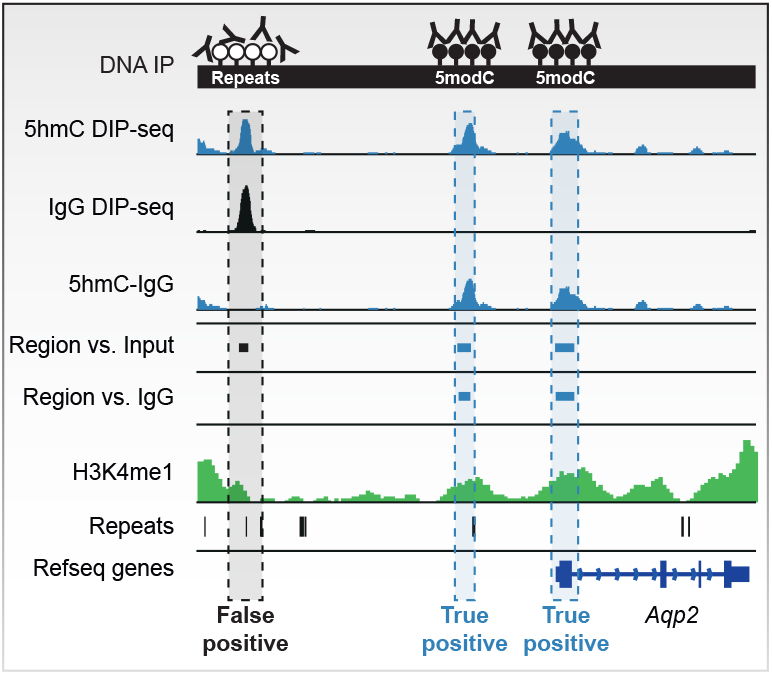
Antibodies in DIP-seq experiments bind repetitive elements which are incorrectly identified as enrichedregions when not controlled for IgG binding.

The prevalence of non-enriched Input DNA as a control in DIP-Seq studies stems from its use in ChIP-seq; Input chromatin helps to control for the different shearing dynamics of closed and open chromatin and for differences in the amplification efficiency of DNA fragments with different base compositions^39^. The preference for Input controls was also fueled by the requirement of a uniform background signal in early peak-calling algorithms^40^. However, Input does not control for non-specific antibody binding. Thus, we strongly suggest that all future DIP-Seq studies perform both an Input and IgG control that will allow for normalization of the effects of sequencing bias and antibody cross-reactivity ((IP-IgG)/Input) ^15, 23^. Indeed, future profiling studies of DNA modifications may be advised to use non-antibody based mapping techniques where possible^41^. Bisulfite sequencing (BS) of 5mC and oxidative BS or TAB-Seq of 5hmC offer quantitative, base-resolution alternatives to (h)MeDIP-seq, but remain prohibitively expensive^42, 43^. The click chemistry based assays, 5hmC-Seal and 5fC-Seal, are low-cost enrichment based techniques that do not exhibit STR enrichment bias, but may be less sensitive than their antibody-based counterparts^16, 20, 24^.

Whereas normalization of DIP-Seq data to an IgG-Seq control represents the optimal approach to generating accurate DIP-Seq profiles, IgG controls are lacking for the majority of published studies. Computational correction of published DIP-Seq data by filtering out sequencing reads containing IgG associated STR motifs is relatively straightforward but is not advised. First, as DNA modifications (5mC, 5hmC, 5fC) do occur at non-CpG dinucleotides in some cell types, complete removal of IgG-STR sequences may result in a loss of biologically significant information^21, 26^ (**Supplementary Fig. 2c, d**). Second, as genomic STR composition differs markedly between species, the set of STRs bound by IgG and the extent of their enrichment is likely to vary in DIP-seq of DNA from different organisms (**Fig. 2e**). Third, as the effect of off-target STR binding increases with decreasing abundance of the target epitope (**Fig. 3a**), *a priori* knowledge of global modification levels in each genome would be required to prevent over-correction of the data. Finally, other experimental variables such as antibody source and sensitivity, DNA denaturation conditions, stringency of washing may also effect the degree of STR-binding observed. Consequently, optimal reanalysis of published DIP-seq data requires the generation of additional IgG-Seq data for each cell type under investigation.

Unexpectedly, we also revealed the potential for contaminating bacterial DNA to confound the results of DIP-Seq studies of trace DNA modifications. The risk of such contaminants has been previously raised with regards to 6mA^10^, which is vanishingly rare in mammals, but highly abundant in many bacterial species that commonly infect mammalian cell cultures, such as *Mycoplasma* and *E.coli.* Fortunately, even minor bacterial contamination of mammalian DNA samples can be identified by comparison of next generation sequencing reads with the genomic sequence of suspected contaminants. Using this approach, we found that up to 15% of reads in published samples of 6mA-DIP-seq in mammals mapped to the *Mycoplasma* genome^31^. Taken together with the results of a recent study that was unable to detect 6mA in mammalian cells using mass spectrometry and our results showing clear enrichment for STRs using the 6mA antibody, a re-evaluation of the extent and origin of 6mA in mammalian studies is advisable^34^. As many bacteria and viruses contain abundant amounts of modified bases including 5mC, 5hmC and 6mA, controlling for contaminating DNA in all DIP-seq assays requires more rigorous application, particularly when the genomic content of target modifications in mammals becomes increasingly rare.

How specific ssDNA molecules become bound to IgG during DNA immunoprecipitation is unclear? It would seem unlikely that the antigen-binding site of antibodies raised against different epitopes (5mC, 5hmC, 5fC, 5caC and 6mA) would all exhibit affinity for the same ssDNA molecules. Alternatively, ssDNA molecules may bind directly to the conserved Fc region of IgG antibodies. Indeed, both ssRNA and ssDNA molecules (‘aptamers’) capable of specifically binding the Fc-region of mouse and rabbit IgG have been reported^44^. Aptamer binding to the Fc regions is highly secondary structure dependent^44^, which may explain enrichment for specific repetitive sequences that have a high probability of forming secondary structures during the DNA denaturation step of DIP. A role for secondary structure in STR-enrichment is further supported by a recent study employing immunoprecipitation of drosophila RNA with a 5hmC antibody (hMeRIP-seq) that found 64% of enriched regions were coincident with AG-rich repetitive sequences^45^. It is tempting to speculate on a function of ssDNA-antibody binding *in vivo* as mechanism to recognize highly structured, single-stranded viral DNA.

Whereas our discovery of unmodified STR binding by IgG has revealed a serious flaw in DIP-seq to date, it will allow the field to minimize the impact of these errors on future DIP based assays and accelerate the discovery of novel findings from the multitude of existing DIP-seq data.

## METHODS

Methods and associated references are available in the online version of the paper.

## ACKNOWLEDGEMENTS

This work was supported by the Swedish Research Council, Åke Wibergs Fund and the LiU-Cancer Network (C.E.N.). R.R.M. and H.J.M. was supported by the Medical Research Council, UK (MC_PC_U127574433).

## AUTHOR CONTRIBUTIONS

C.L., S.V., K.D., and H.K.M. performed experiments, A.L., C.E.N. and S.V. analyzed data, A.L. R.R.M. and C.E.N. wrote the manuscript and H.V., H.G., R.R.M., M.B. and C.E.N. supervised the work.

## SUPPLEMETARY FIGURE LEGENDS

**Figure S1.**
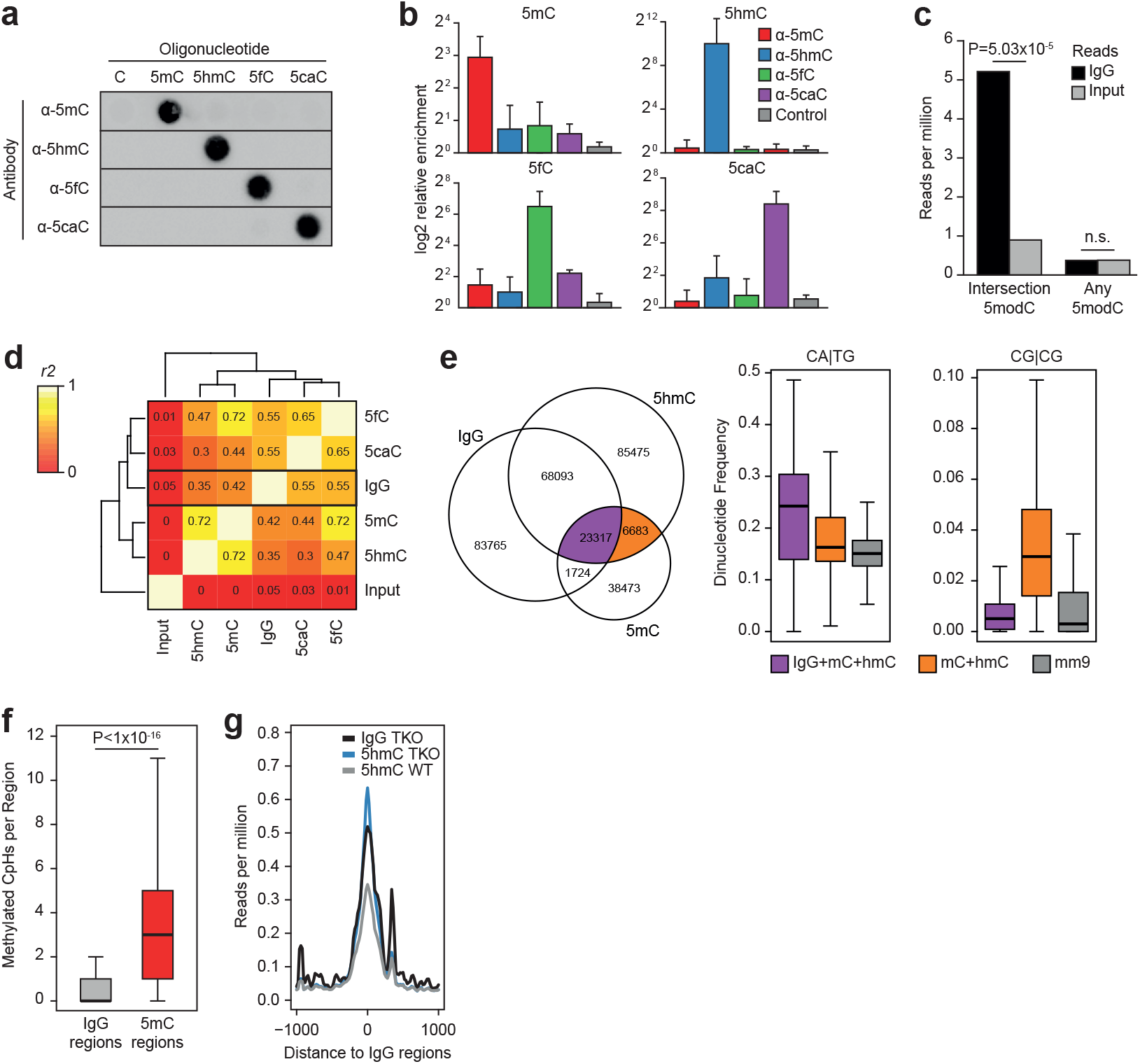
(**a, b**)Immuno dot blot (**a**) and ELISA (**b**) of 5mC, 5hmC, 5fC and 5caC antibodies showing specificity for their respective mark in synthetic 426 bp oligos containing the different marks. (**c**) Enrichment of IgG or Input reads over the intersection of 5modC (5mC+5hmC+5fC+5caC) enriched regions or non-intersecting (5mC/5hmC/5fC/5caC) regions. P-values calculated using T-test. (**d**) Correlation matrix of number of enriched DIP-seq regions per Mbp of mm9. All 5modC marks show correlation with IgG DIP-seq compared to Input. Correlation was calculated as Pearson correlation. (**e**) Venn diagram of overlapping enriched regions for 5hmC, 5mC and IgG (left). Dinucleotide frequencies for overlapping regions, boxplots represent median and top/bottom quartiles (right). (**f**) Number of methylated CpH from WGBS data per enriched IgG or 5mC region. Boxplots represent median and top/bottom quartiles, P-values calculated using Mann-Whitney U-test. (**g**) Enrichment profile of IgG and 5hmC in TKO and WT mESCs over regions enriched for IgG showing highly similar profiles.

**Figure S2.**
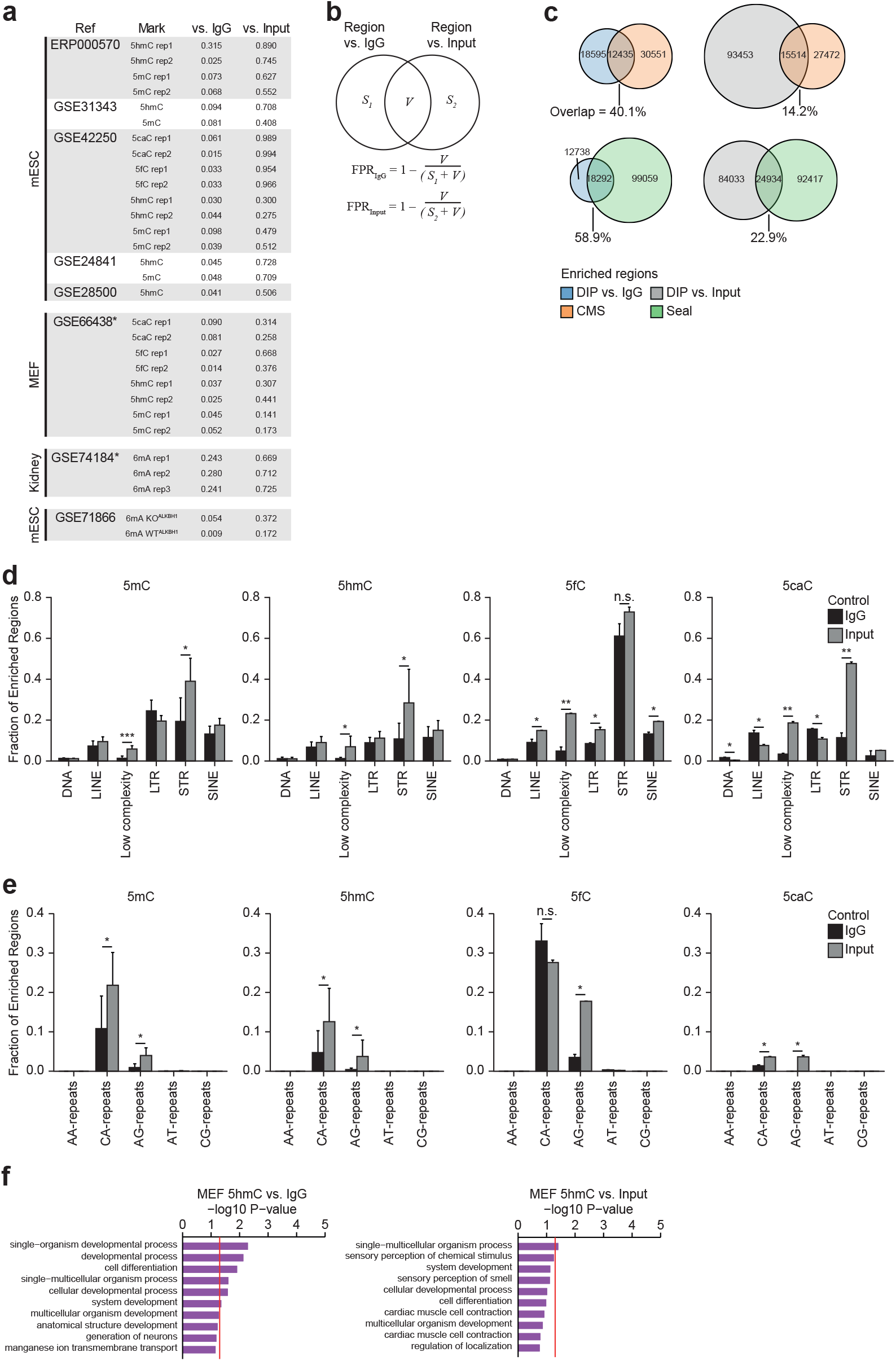
(**a**) Estimated false positive rate for individual mESC or MEF datasets. ^*^Estimated based on controls from mESCs. (**b**) False positive rate (FPR) was estimated based on the inverse fraction of regions identified by both Input and IgG versus total regions. (**c**) Venn diagram of enriched 5hmC regions in mESCs with different techniques showing higher overlap for 5hmC DIP-seq when using IgG controls. (**d, e**) Fraction of enriched 5modC regions identified using IgG or Input overlapping repetitive elements (**d**) and dinucleotide repeats (**e**). Presented as mean ± s.d of 2–7 biological replicates. *P<0.05, **P<0.01, ***P<0.001, T-test. (**f**) GO term enrichment of biological processes for 5hmC enriched regions in mouse embryonic fibroblasts (MEFs)Lentini *et al.*using either IgG or Input controls. Stronger enrichment for developmental terms is observed when using the IgG controls.

## SUPPLEMNTARY METHODS

### Cell culture

J1 mouse embryonic stem cells (mESCs; WT, male) were originally derived from the 129S4/SvJae strain. TKO (Dnmt1-/-, Dnmt3a-/-, Dnmt3b-/-) mESCs were derived from J1 mESCs^46^. Both cell lines were cultured in a humidified incubator at 5% CO_2_, 37°C on 0.2% gelatin coated tissue culture plastic in DMEM (Dulbecco’s modified eagle medium) supplemented with 15 % fetal calf serum, 0.1 mM non-essential amino acids (Sigma-Aldrich, MI, USA), 1 mM sodium Pyruvate (Sigma-Aldrich, MI, USA), 1 % Penicillin/Streptomycin, 2 mM L-glutamine, 0.1 mM beta-mercaptoethanol (Thermo Fisher, CA, USA), and ESGRO LIF (Millipore, MA, USA) at 500U/mL. mESCs were passaged every 2-3 days using trypsin/EDTA.

### DNA extraction

Snap frozen cell pellets were treated with RNAse cocktail (Ambion, CA, USA) for 1 hour at 37°C followed by proteinase K treatment overnight at 55°C. DNA was extracted by standard phenol chloroform/ethanol precipitation and eluted in TE.

### DIP-qPCR

1.5 μg genomic DNA was sonicated to fragments ranging between 100-1000 bp with a peak at 400 bp using a BioRuptor (Diagenode, Belgium), denatured at 95°C for 10 min then cooled on wet ice for 10 min. 10% of samples were saved as Input and the remaining DNA was resuspended in 10x IP buffer (10 mM Na-Phosphate (mono-dibasic), 1% NaCl, 0.05% Triton X–100, pH 7.0). Immunoprecipitations were performed using 1μg anti-5hmC antibody (Active Motif, #39769) for 12h at 4°C using constant rotation. Protein G dynabeads (Invitrogen, CA,USA, #100–03D) were washed twice in 0.1% PBS-BSA then added to the IP mixture for 1h at 4° using constant rotation. Beads were washed three times for 10 min using cold 1x IP buffer then resuspended in digestion buffer and incubated with 8 U Proteinase K (New England Biolabs, MA, USA) for 2h 1.5h at 50°C, 800rpm in 50 mM Tris,10 mM EDTA0.5% SDS, pH 8.0 and purified using DNA Clean & Concentrator^™^-5 kit (Zymo Research, USA). Quantitative PCR was performed on a 7900HT real-time cycler (Applied Biosystems,CA, USA) using SYBR green master mix (Applied Biosystems, CA, USA). qPCR primers use **Supplementary Table 4**. hMeDIP qPCR primer sequences are listed in **Supplementary Table 4**, below.

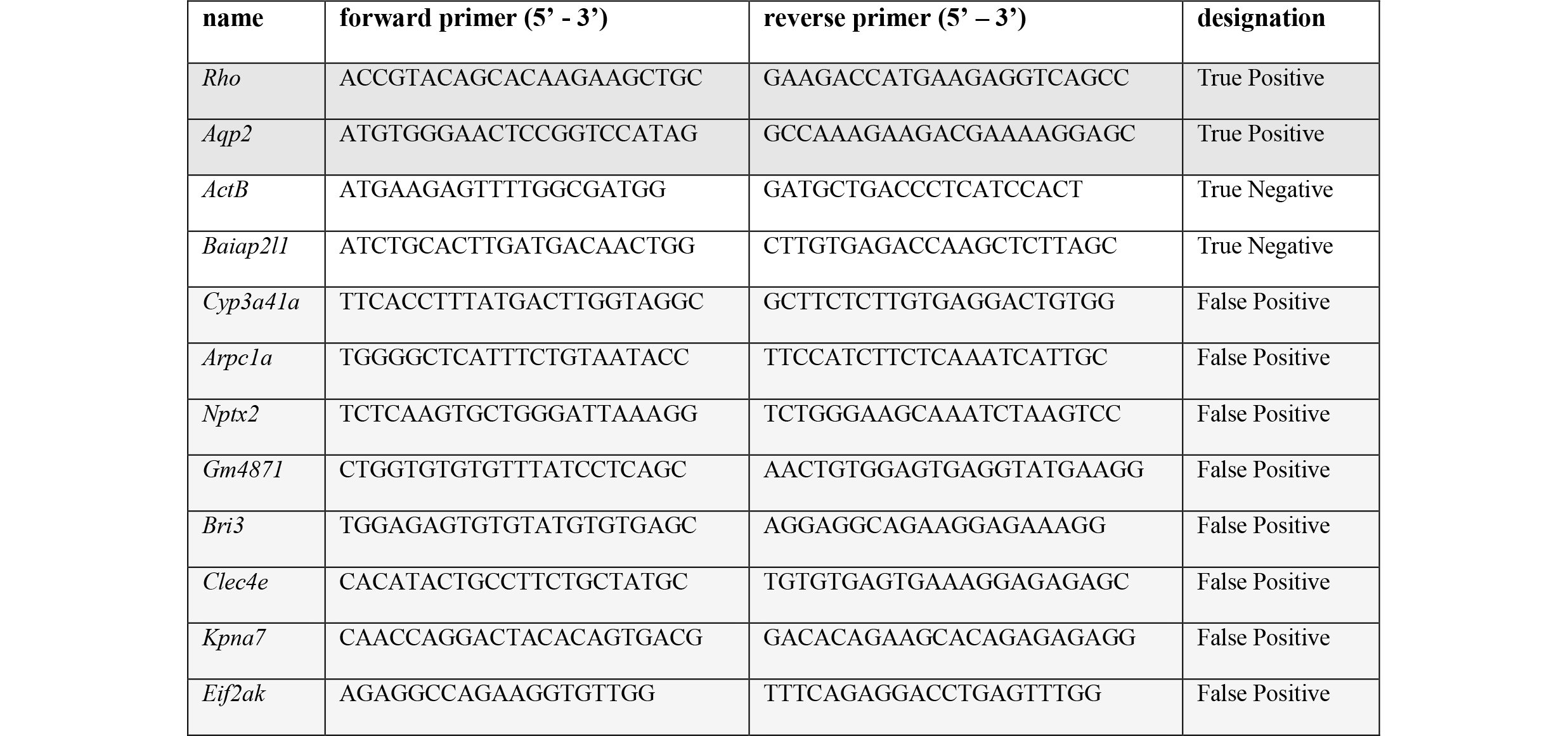

### Quantification of cytosine modifications using mass spectrometry

1 μg of DNA was heat denatured at 100 °C for 5 min in 20μL H2O then immediately cooled on ice. 10 μl P1 Nuclease (0.02 U/μl in 90 mM AmAc, 0.3 mM ZnSO4, pH 5.3) was added followed by incubation at 50 °C for 2 h. 10 μl Alkaline phosphatase (0.08 U/μl in 200 mM TRIS-HCl, 0.40 mM EDTA, pH 8) was added followed by incubation at 37 °C for 30 min. Proteins were precipitated by the addition of 160 μl cold acetonitrile. Following centrifugation at 17000 x g for 5 min, 180 μl of the supernatant was evaporated under nitrogen and reconstituted in 40 μl 0.1% formic acid. The chromatographic system consisted of an Acquity UPLC (Waters, MA, USA) and a Xevo triple quadrupole mass spectrometer (Waters, MA, USA). The extracts were separated on an HSS T3 column (150x2.1 mm, 1.7 μm, Waters, MA, USA) at 45°C and a flow rate of 450 μl/min using a gradient elution with 0.05% acetic acid and methanol, 0–1.3 min 2% B; 1.3–5.5 min 2–9% B; 5.5–7.5 min re-equilibration at 2% B. For dC a 1 μl injection was made and for mC, hmC, fC and caC a 15 μl injection was made. Analytes were detected in the multi reaction monitoring (MRM) mode using three time windows with the following transistions 0–2.3 min – C (228->95 & 228->112) and hmC (258->124 & 258->142); 2.3–4 min – mC (242->109,242->126) and caC(272->138, 272->156); 4-7.5 min – fC (256->97, 256->140).

### Immuno dot-blot

10 ng 426 bp oligos containing 5mC, 5hmC, 5fC, 5caC or C (GeneTex, CA, USA) was denatured at 95°C for 15 min in 0.4M NaOH and 10mM EDTA then immediately cooled on ice. Samples were applied to a positively charged nylon membrane under vacuum using a Dot Blot Hybridisation Manifold (Harvard Apparatus, MA, USA). The membranes were briefly washed in 2X SSC buffer (0.3M NaCl, 30mM NaCitrate) then crosslinked using a UV Stratalinker 1800 (Stratagene, CA, USA) and baked at 80°C for 2 hours. Membranes were blocked in casein blocking buffer (Li-Cor) for 15 min at 4°C then incubated with an antibody against 5mC (1:3000, Zymo #A3001), 5hmC (1:3000, ActiveMotif #39791), 5fC (1:3000, ActiveMotif #61227) or 5caC (1:3000, ActiveMotif #61229) for 1h at 4°C. Membranes were washed 3 times for 5 min in TBS-Tween (0.05%) then incubated with a HRP conjugated goat-anti-rabbit antibody for 5hmC, 5fC and 5caC (1:3000, Bio-Rad #1706515) or goat-anti-mouse for 5mC (1:3000, Bio-Rad #1706516). Following treatment with Clarity Western ECL substrate (Bio-rad, CA, USA), membranes were scanned on a ChemiDoc MP imaging system (Bio-Rad, CA, USA).

### ELISA

426 bp dsDNA oligos containing 5mC, 5hmC, 5fC, 5cacC or C (GeneTex, CA, USA) was diluted to a concentration of 50ng/mL in coating buffer (1M NaCI, 50 Mm Na2PO4, 0.02% (w/v) NaN3, pH 7.0) then 50μl were placed into each well of black 96-well plates (4titude, UK) and incubated overnight at 37°C. Plates were blocked for 1h at room temperature in Blocker Casein in PBS (Thermofischer Scientific, MA, US) followed by washing with 100 μl PBS containing 0.1% (v/v) Tween 20. Wells were incubated with 50μl of their respective antibodies (1:1000, see above) for 1h at room temperature, then washed 3 times and incubated with 50μl of horseradish peroxidase (HRP)-conjugated goat-anti-mouse or goat-anti-rabbit antibody (1:5000, see above) for 30 min. Plates were treated with 70μl of Clarity Western ECL substrate (Bio-rad, CA, USA) for 5 min then scanned in a Spark 10M multimode microplate reader (Tecan Trading AG, Switzerland).

### Uniform analysis pipeline for processing of published DIP-Seq data

Raw 5modC DIP-seq sequencing data was downloaded from GSE42250, GSE24841, GSE31343, ERP000570, GSE28500 and aligned to the mouse genome (mm9) using Bowtie2^47^ (bowtie2-N 1-L 30). Genomic coverage was calculated using Bedtools^48^ (bedtools genomecov -bg -split) then normalized as reads per million mapped (RPM) for visualization. Identification of enriched regions was performed using MACS2^49^ (macs2 --bw=200 -p 1e-5) using IgG or Input controls from the same study where possible otherwise IgG or Input samples from the above studies were pooled and randomly subsampled to 20 million reads as controls. Unless otherwise stated, 5modC enriched regions were identified using IgG controls and IgG enriched regions using Input.

6mA DIP-seq data was downloaded from GSE71866, GSE74184 and GSE76740 and processed as 5modC DIP-seq data (see above).

Bisulfite sequencing data was obtained from GSE41923 and aligned to a bisulfite converted mm9 index using Bismark^50^ (bismark −N 1). Methylation levels of Cytosines in both CpG and non-CpG contexts were extracted (bismark_methylation_extractor) and bases with at least 5X coverage were used for analysis.

Raw 5hmC-Seal data was downloaded from GSE41545 and processed as DIP-seq data (see above) and anti-CMS was downloaded from GSE28682 and aligned using Bismark^50^ with the same settings as for DIP-seq (bismark -N 1 -L 30).

TET1 ChIP-seq data was downloaded from GSE24843 and histone ChIP-seq data for mESCs was obtained from the ENCODE project^51^ and processed as DIP-seq data (see above).

See **Supplementary Table 1** for specification of files used for each analysis/figure.

### Estimation of falsely enriched regions

Enriched regions were obtained from MACS2 using either pooled IgG or Input from mESCs as control (see above). True regions were considered when an overlapping region was called for both IgG and Input controls and falsely enriched regions were calculated as the inverse fraction for either control (**Supplementary Fig. 2b**).

### Motif enrichment of FASTQ files

FASTQ files were trimmed of adapters using ea-utils^52^ (fastq-mcf -x 0 -q 0 -k 0 -s 4.6) then randomly subsampled to 1 million reads and subjected to *de novo* motif enrichment analysis using Homer2^53^ (homer2 denovo -len 12). Input samples from the same study was used as background when available, otherwise a pooled input from multiple studies was used (see above). Correlation between motif PWMs was performed using Pearson correlation as implemented in TFBStools^54^ (PWMsimilarity), subject motifs were repeated once to account for base shifts. To identify if identified motifs belong to a certain repeat class, motif PWMs were mapped to repeats in mouse (RepBase v22.01^55^) using Homer2^53^ (scanMotifGenomeWide.pl).

### Taxonomic annotation of sequence reads

Taxonomic annotation of raw sequencing reads was performed by aligning reads to a custom reference genome of mm9 combined with the bacterial genomes of Mycoplasma species M.arginini (ASM154797v1), M.hyorhinis_dbs1050 (ASM49681v1), M.hyorhinis_gdl1 (ASM24112v1), M.hyorhinis_hub1 (ASM14570v1), M.hyorhinis_mcld (ASM21129v1), M.hyorhinis_sk76 (ASM31363v1) as well as E.coli (HUSEC2011CHR1) using Bowtie2^47^, with the same settings as outlined above. For determination of short tandem repeat (STR) fraction of species genomes, Tandem Repeat Finder^56^ (TRF) results for genomes (ce10, danRer10, dm6, hg38, mm10) was obtained from UCSC. For *Xenopus laevis*, the genomic sequence was obtained from Xenbase (Xenla9.1) and STRs was identified using TRF 4.09 with recommended settings and a maximum period size of 12 (trf 2 7 7 80 10 50 12).

### GO term enrichment analysis

Top 500 enriched regions were mapped to the nearest gene within 10kb and enrichment of GO terms biological processes was performed using PANTHER^57^ with default settings.

### Statistical analysis

All statistical analysis was performed using the statistical programming language R^58^ unless otherwise stated. P-values <0.05 were considered significant.

